# A Ubiquitin-Binding Domain that Does Not Bind Ubiquitin

**DOI:** 10.1101/375832

**Authors:** Michael Lim, Joseph A. Newman, Hannah L. Williams, Hazel Aitkenhead, Opher Gileadi, Jesper Q. Svejstrup

## Abstract

Ubiquitylation, the post-translational linkage of ubiquitin moieties to lysines in target proteins, helps regulate a myriad of biological processes. Ubiquitin, and sometimes ubiquitin-homology domains, are recognized by ubiquitin-binding domains, including CUE domains. CUE domains are thus generally thought to function exclusively by mediating interactions with ubiquitylated proteins. The chromatin remodeler, SMARCAD1, interacts with KAP1, a transcriptional corepressor. We show that the SMARCAD1-KAP1 interaction is direct and involves the first SMARCAD1 CUE domain (CUE1) and the RBCC domain of KAP1. A structural model of the minimal KAP1 RBCC-SMARCAD1 CUE1 complex based on X-ray crystallography analysis is presented. Remarkably, the CUE1 domain, which resembles a canonical CUE domain, recognizes 2 clusters of exposed hydrophobic residues on KAP1, but these are presented in the context of a coiled-coil domain, not in a structure resembling ubiquitin. Together, these data challenge the well-established dogma that CUE domains exclusively recognize the ubiquitin-fold.

## Introduction

CUE domains are ubiquitin-binding domains (UBDs) that interact with ubiquitin by occupying the hydrophobic pocket centred on the highly conserved ubiquitin I44 residue^1–3^. As is typical of most UBD-ubiquitin interactions, CUE-ubiquitin interactions are relatively weak, reflecting a comparatively small interaction surface of only ~400Å^2 4–7^. CUE domains have two main conserved sequence elements – a methionine-phenylalanine-proline (MFP) motif and a di-leucine (LL) repeat, both of which are essential for ubiquitin binding^4,5,8^. With the rare exception of ubiquitin-homology domains (UbHs), ubiquitin remains the only known ligand of CUE domains^1,9^. As UBDs are essential in detecting the ubiquitylation status of their partner proteins, they often play a crucial role in mediating the regulation of biological processes by ubiquitylation. Thus, mutation of the UBD of a protein to perturb its interaction with ubiquitin is often a reasonable starting point for interrogating the biological function of that protein.

SMARCAD1 is a candidate for investigation in this manner, as it has a pair of CUE domains and biological functions that merit further characterization at a mechanistic level. SMARCAD1 is a chromatin remodeler, a member of the SWI2/SNF2-like family of enzymes that couple the energy released from ATP hydrolysis to repositioning, ejecting, or restructuring nucleosomes^10^. SMARCAD1 has homologues spanning considerable evolutionary distance – its homologues are Fun30 in budding yeast, Fft1, Fft2 and Fft3 in fission yeast, and Etl1 or HEL1 in mouse^11,12^. By virtue of its split-ATPase domain, SMARCAD1 is most closely related to the Swr1-like group of remodelers, including INO80 and SWR1^11^. Specific biochemical activity has not, to date, been demonstrated for human SMARCAD1. However, the INO80 and SWR1 remodelers primarily facilitate histone dimer exchange, and the SMARCAD1 homologue Fun30 similarly preferentially mediates histone exchange over nucleosome sliding^13–15^. It seems likely that SMARCAD1 will play a comparable role in mediating histone exchange in vertebrates.

Functionally, SMARCAD1 has been implicated in facilitating homologous recombination by promoting end-resection, and in maintaining constitutive heterochromatin through DNA replication^16,17^. Its *S. pombe* homologue, Fft3, has been reported to modulate nucleosome exchange – both promoting it, in association with elongating RNA polymerase II to relieve the nucleosomal barrier to transcription, and suppressing it, to facilitate heterochromatin maintenance^18,19^. In a Xenopus egg extract system, SMARCAD1 promotes Msh2-dependent mismatch repair on a chromatin template^20^. Collectively, these observations link SMARCAD1 to the regulation of various processes in the context of the chromatin environment^16–24^.

KAP1 (also known as TRIM28 and TIF1β) is the major interaction partner of SMARCAD1, with the two proteins forming a tight complex^16,25^. KAP1 is ubiquitously expressed, and is implicated in transcription repression, heterochromatin formation, and in DNA repair, amongst other functions^26–30^. KAP1, in common with other TRIM proteins, has a distinctive RBCC domain – a composite of <u>R</u>ING, tandem <u>B</u>-box and <u>c</u>oiled-<u>c</u>oil domains. Additional features of KAP1 are a central HP1-box and a C-terminal PHD-bromodomain^26,27^.

Recent structural work has demonstrated that the RBCC domain of the homologous TRIM25 protein forms an elongated structure, dominated by its coiled-coil domain homo-dimerizing in an antiparallel manner^31^. Specifically, each coiled-coil subunit forms a hairpin, with a short arm consisting of a C-terminal linker region that folds back onto a long arm, comprised of the coiled-coil domain proper. The coiled-coil has a distinctive palindromic arrangement (i.e. 7-7-7-7-11-11-11-11-7-7-7-7) of heptad (denoted ‘7’) and hendecad (denoted ‘11’) repeats^31^. The centrally located hendecad repeats form an underwound R-handed helix that is infiltrated by the linker helices of the short arm, resulting in a central four-helical bundle. The N-terminal RING and B-boxes are located at the apices of the coiled-coil, whereas C-terminal domains (e.g. the PHD-bromodomain in KAP1) presumably protrude from either side of the centre of the coiled-coil^31^. Other TRIM proteins share this overall structural configuration^31–33^. This architectural organization enables TRIM proteins to assemble protein domains in a modular fashion; hence, TRIM proteins should excel as scaffolds for recruiting specific cellular machinery to target locations. It is also conceivable that the considerable exposed surface created by TRIM coiled-coil dimerization might represent yet another interaction surface.

Given the functional significance of the SMARCAD1 protein, and the undetermined significance of its CUE domains, we focused on the SMARCAD1 CUE domains for further investigation.

## Results

### SMARCAD1 and KAP1 Interact Directly in a Ubiquitylation-Independent Manner

The SMARCAD1 protein contains a pair of tandem CUE domains of largely unknown function (Figure 1A). We generated human cell lines, depleted of endogenous SMARCAD1 by shRNA knockdown, which were then reconstituted with shRNA-resistant cDNA encoding either FLAG-tagged wild type SMARCAD1 or SMARCAD1 with point mutations in the CUE domains (“CUE1mt,2mt”), under the control of a doxycycline-inducible promoter (Figure 1B). The CUE1mt,2mt protein should be compromised in its ubiquitin-binding ability, as each of its CUE domains possesses four alanine substitutions targeting the conserved, hydrophobic MFP and di-leucine motifs that are integral to the ubiquitin interaction surface^2^. As expected, KAP1 co-immunoprecipitated with wild type SMARCAD1 (Figure 1C, lane 8). Strikingly, however, point mutation of the tandem CUE domains abrogated the interaction (lane 9). As CUE domains are ubiquitin-binding domains, SMARCAD1-KAP1 interaction might involve ubiquitylated KAP1. We explored whether cells possess a pool of ubiquitylated KAP1 by isolating ubiquitylated proteins from 293T human cell extract using MultiDsk affinity purification^34^, and then examining KAP1 ubiquitylation by Western blot analysis. Surprisingly, however, no ubiquitylated KAP1 was detected by this approach (data not shown).

**Figure 1.**
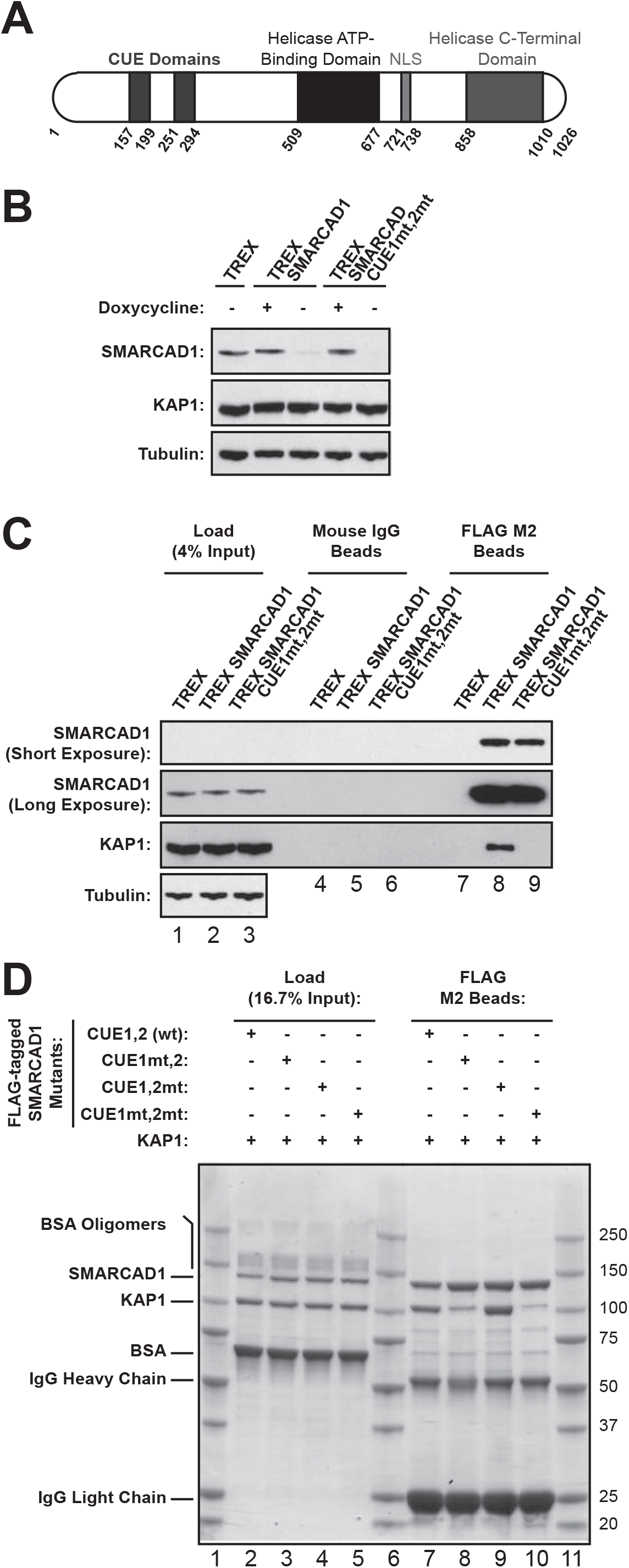
The SMARCAD1-KAP1 interaction depends on the first SMARCAD1 CUE domain. **A**. Domain architecture of the human SMARCAD1 protein, with amino acid positions listed below. **B**. Stable HEK293 T-REx cell lines, depleted of endogenous SMARCAD1, but inducibly (i.e. in the presence of doxycycline) expressing exogenous FLAG-tagged SMARCAD1 at approximately normal levels. The cells were reconstituted with either wild type protein, or SMARCAD1 with inactivating mutations in both CUE domains (“CUE1mt,2mt”). **C**. Wild type and SMARCAD1 CUE1mt,2mt was enriched by FLAG immunoprecipitation. Only wild type specifically co-immunoprecipitates KAP1 (compare lanes 8 and 9). **D**. The SMARCAD1-KAP1 interaction reconstituted with non-ubiquitylated, purified proteins, expressed in *E. coli*. Mutation of SMARCAD1 CUE1 significantly compromises KAP1 binding (compare lanes 7 and 8).

We previously found that the CUE domain of the yeast Def1 protein recognizes the ubiquitin-homology (UbH) domain of Elongin A (Ela1)^9^. Thus, the possibility that KAP1 similarly interacts directly with SMARCAD1 in a ubiquitylation-<u>in</u>dependent manner was considered. Exploiting the inability of prokaryotes to ubiquitylate proteins, recombinant SMARCAD1 and KAP1 were purified following expression in *E. coli* (Figure S1A and D), and then tested for their ability to interact *in vitro*. After mixing, KAP1 indeed co-immunoprecipitated with SMARCAD1 (Figure 1D, lane 7), indicating that these proteins interact directly, in a ubiquitylation-independent fashion. Remarkably, the purified SMARCAD1 CUE1mt,2mt protein showed little binding to KAP1 (lane 10), recapitulating the observation in mammalian cells. Similarly, SMARCAD1 protein with point-mutations only in the first CUE domain (“CUE1mt,2”) was significantly compromised in its ability to bind KAP1 (lane 8), while point mutation of only the second CUE domain (“CUE1,2mt”) did not perceptibility affect the interaction (lane 9). Collectively, these data indicate that the SMARCAD1-KAP1 interaction is a direct protein-protein interaction that is mediated via the first CUE domain of SMARCAD1, and that the tandem CUE domains of SMARCAD1 are not functionally redundant for this function.

We next investigated whether a stable SMARCAD1-KAP1 complex could be reconstituted *in vitro*. A simple strategy of sequential FLAG- and HA-affinity purification from a mixture of FLAG-tagged SMARCAD1 and HA-tagged KAP1 proteins was adopted (Figure S2A). It was, indeed, possible to reconstitute a SMARCAD1-KAP1 complex containing stoichiometric quantities of each of the partner proteins (Figure S2B, lane 8). Furthermore, the reconstituted SMARCAD1-KAP1 complex behaved as expected for a stable protein complex on gel filtration chromatography (Figure S2C). We conclude that SNARCAD1 and KAP1 not only interact, but form a highly stable protein complex.

### The RBCC Domain of KAP1 and the First CUE Domain of SMARCAD1 are Necessary and Sufficient for the SMARCAD1-KAP1 Interaction

Close inspection of the domain architecture of KAP1 did not offer any indications about the region of the protein that is recognized by the first CUE domain of SMARCAD1. Consequently, limited tryptic digestion was used to identify KAP1 fragments that reflect the tertiary structure of the protein – three KAP1 fragments were relatively resistant to tryptic digestion (Figure 2A, lane 5). To map these fragments, the sequences of their N-termini were determined by Edman degradation, while their mass was measured to a high level of accuracy by intact molecular weight mass spectrometry. Interpreted in light of the knowledge that trypsin cleaves after lysine or arginine residues, the largest KAP1 fragment (“Fragment 1”, S33-K434, 45kDa) was found to span the RBCC domain of KAP1 (Figure 2B). Further cleavage yields a second fragment, which encompasses the second B-box and the coiled-coil domain (“Fragment 2”, S200/D202-K434, 27kDa). The final fragment (“Fragment 3”, L592-P835, 26kDa) spans the C-terminal PHD-bromodomain (Figure 2B).

**Figure 2.**
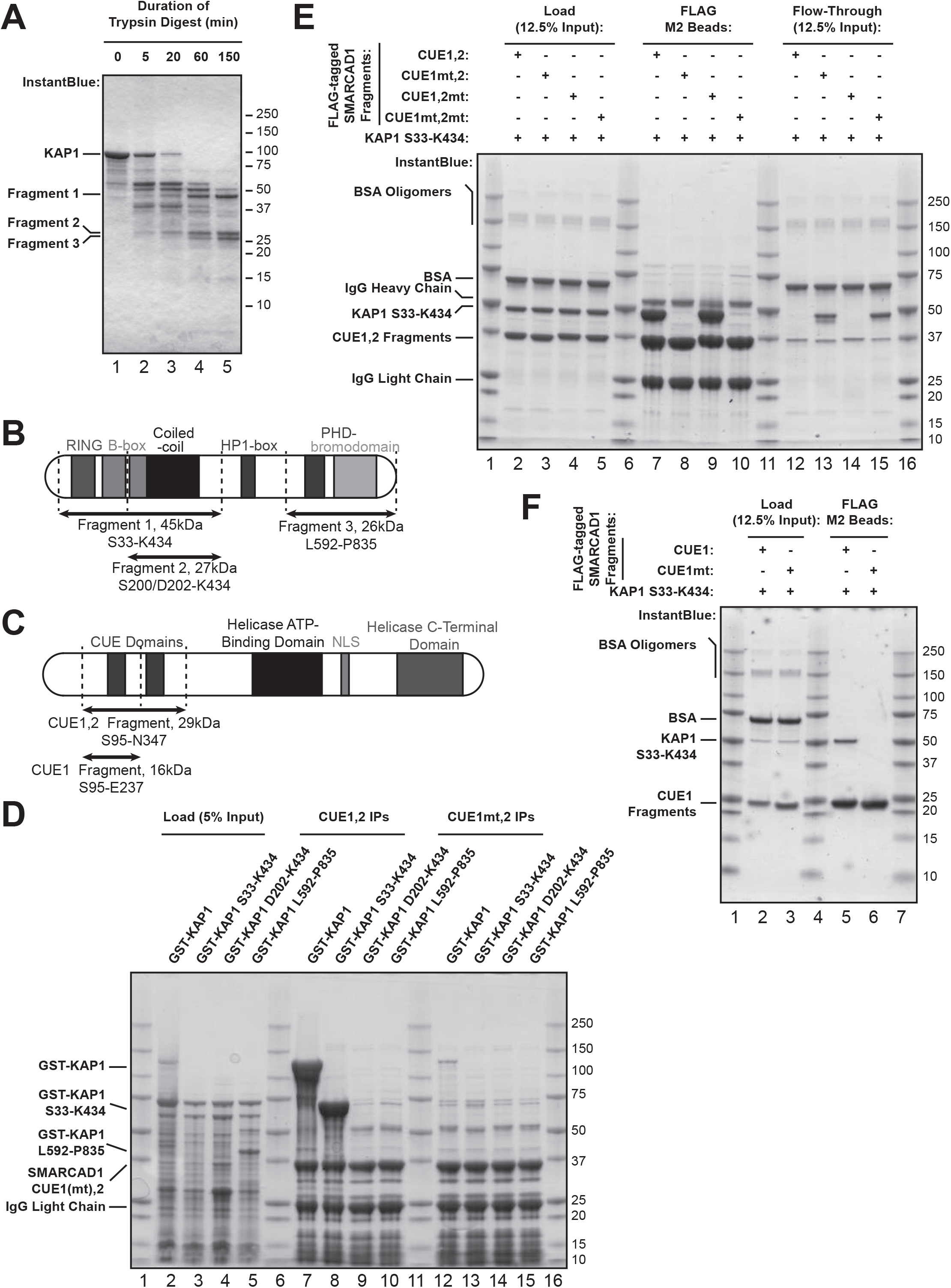
Defining the minimal requirements for the SMARCAD1-KAP1 interaction. **A**. Limited tryptic proteolysis of purified recombinant KAP1 yields three main fragments that are relatively resistant to trypsin. **B**. The three trypsin-resistant KAP1 fragments mapped by Edman degradation and intact molecular weight mass spectrometry. The domains encompassed by each of these fragments are depicted. **C**. Schematic representation of SMARCAD1 fragments spanning either both CUE domains (“CUE1,2”) or only the first CUE domain (“CUE1”) that were expressed and purified. **D**. Immobilized SMARCAD1 CUE1,2 fragment enriches for both full-length KAP1 (lane 7) and KAP1 S33-K434 (i.e. fragment 1, lane 8), which spans the entire RBCC domain, from *E. coli* extract. Binding is specific to the wild type CUE1,2 fragment; point mutation of the first CUE domain (“CUE1mt,2”) abrogates the interaction (lanes 12 and 13). **E**. SMARCAD1-KAP1 interaction recapitulated *in vitro* by the SMARCAD1 CUE1,2 fragment and KAP1 S33-K434 (lane 7). CUE1,2 with inactivating mutations in the first CUE domain are unable to bind to KAP1 S33-K434 (lanes 8 and 10). **F**. The first CUE domain of SMARCAD1 and the KAP1 RBCC domain (S33-K434) are necessary and sufficient for the SMARCAD1-KAP1 interaction.

The three trypsin-resistant fragments were expressed in *E. coli* as GST fusion proteins. Crude bacterial protein extracts containing these GST fusion proteins, or full-length GST-KAP1 as a control, were then incubated with beads immobilised with a SMARCAD1 CUE1,2 fragment (S95-N347), encompassing both of the CUE domains (Figure 2C). The SMARCAD1 CUE1,2 fragment strongly enriched both full-length KAP1 and the S33-K434 RBCC fragment (Figure 2D, lanes 7 and 8). Crucially, this interaction depended on the integrity of the first CUE domain of SMARCAD1: point mutation in this domain completely abrogated binding (lanes 12 and 13). It is worth noting that KAP1 D202-K434 was not readily expressed in *E. coli*, exhibited poor solubility, and frequently precipitated during purification (data not shown). Thus, the apparent inability of the D202-K434 fragment to bind to the SMARCAD1 CUE1,2 fragment (lanes 9 and 14) might potentially simply reflect its tendency to aggregate and low abundance in the extracts.

To further confirm that KAP1 S33-K434 is responsible for the interaction with SMARCAD1, it was purified and tested in a binding assay similar to that described above. Four different FLAG-tagged SMARCAD1 CUE1,2 fragments – the wild type fragment and those resulting from mutations targeting an individual or both CUE domain(s) – were assessed for their ability to bind the KAP1 fragment. Stoichiometric quantities of KAP1 S33-K434 bound to either the wild type CUE1,2 fragment, or the CUE1,2mt mutant (i.e. inactive second CUE domain) (Figure 2E, lanes 7 and 9). Consistent with what was observed with the full-length proteins, KAP1 S33-K434 was incapable of interacting with either the CUE1mt,2 mutant (i.e. inactive first CUE domain), or the CUE1mt,2mt double mutant (lanes 8 and 10). Moreover, a purified wild type CUE1 fragment (S95-E237), but not the CUE1mt mutant version of it, bound purified KAP1 S33-K434 (Figure 2F, compare lanes 5 and 6), further confirming that the second CUE domain of SMARCAD1 is dispensable for SMARCAD1-KAP1 interaction.

Taken together, these data demonstrate that the first CUE domain of SMARCAD1 and the RBCC domain of KAP1 (S33-K434) are necessary and sufficient to mediate a direct interaction between SMARCAD1 and KAP1.

### Crystal structure of the KAP1 RBCC-SMARCAD1 CUE1,2 Complex

CUE domains recognize ubiquitin and, occasionally, UbH domains. The direct, CUE domain-dependent interaction between SMARCAD1 and non-ubiquitylated KAP1 thus raised the possibility that KAP1 harbours a UbH domain. If that hypothesis were true, we reasoned that an excess of ubiquitin might be able to interfere with the SMARCAD1-KAP1 interaction. Strikingly, however, even a hundred-fold (Figure S3A, lane 10) or a thousand-fold (data not shown) molar excess of mono-ubiquitin failed to affect the ability of the SMARCAD1 CUE1,2 fragment to interact with KAP1 S33-K434. In apparent agreement with these data, there is no convincing sequence homology between ubiquitin and the KAP1 RBCC domain. While not conclusive, these data provided the first arguments against the presence of a simple UbH domain in KAP1 whose binding to SMARCAD1 could be competed with ubiquitin.

To further investigate the unprecedented possibility of a CUE domain binding a ligand structurally distinct from ubiquitin, we crystallized the KAP1 RBCC domain (residues 53 to 434) in complex with the SMARCAD1 CUE1,2 fragment (residues 94-347). The structure was determined to 5.5Å resolution by single-wavelength anomalous diffraction, using the intrinsic anomalous signal of the zinc ions (i.e. 4 per KAP1 subunit). Despite the relatively low resolution, the experimental phases were of excellent quality (Figure 3A) and were significantly improved by solvent flattening algorithms due to the extremely high solvent content of 92%, which is amongst the highest in the PDB archive. The high solvent content reflects the unusual packing arrangement within the crystal, which has a unit cell of 300Å diameter in the form of a large proteinaceous cage with internal voids (Figure S4A). Despite numerous permutations of constructs and crystallization conditions, this was the highest resolution achieved. It was, nevertheless, possible to build a full atomic model of the complex, aided by the availability of templates for individual domains, and the ability to determine the sequence register unequivocally at sites of zinc ion coordination and at breaks in secondary structural elements.

**Figure 3.**
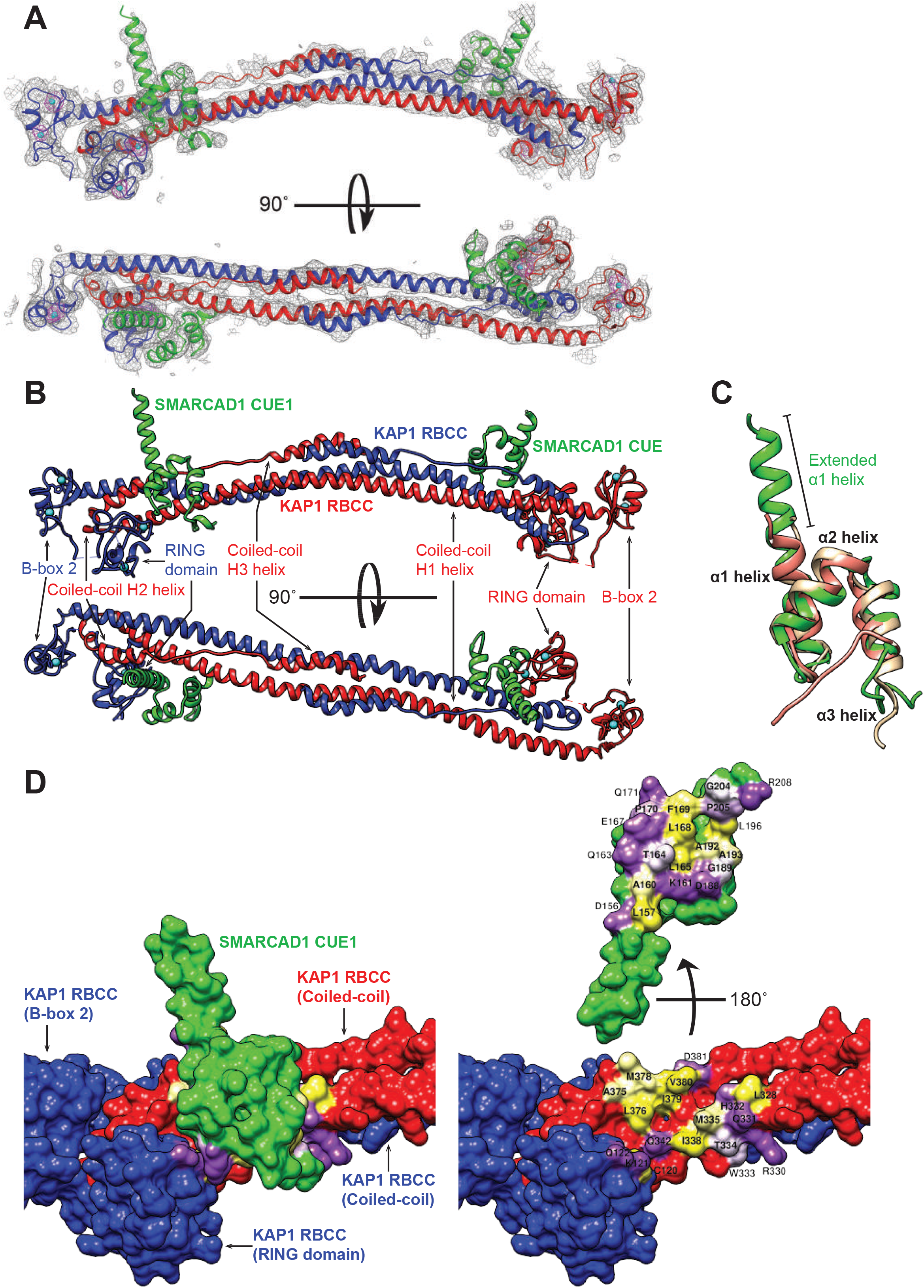
A structural model of the KAP1 RBCC-SMARCAD1 CUE1,2 complex. **A**. Electron density map of the KAP1 RBCC-SMARCAD1 CUE1,2 complex. One KAP1 RBBC chain is coloured red, and the other blue; the CUE1 domains are coloured green. Electron densities corresponding to the first B-box of KAP1 and the second SMARCAD1 CUE domain were not observed. **B**. KAP1 RBCC homo-dimerizes, adopting a barbell-like appearance. The coiled-coil domains mediate homo-dimerization in an anti-parallel fashion and form the elongated central helical structure. The RING and B-box domains are located at either end of the coiled-coil helix. A SMARCAD1 CUE1 domain binds to each end of the KAP1 RBCC dimer. The KAP1 interaction interface is primarily comprised of an exposed surface of the coiled-coil domain of one RBCC subunit, with minor contributions from the RING domain of the other RBCC subunit. The domains are coloured as in Figure 3A. **C**. The SMARCAD1 CUE1 domain (green) resembles canonical CUE domains. The CUE domains of CUE2p (tan) and gp78 (salmon) are superimposed on top. **D**. The SMARCAD1 CUE1-KAP1 RBCC interaction surface. In the figure on the right, the CUE1 domain has been rotated 180° off KAP1. Residues involved in the interaction are labelled and coloured according to their hydrophobicity: yellow = hydrophobic, white = hydroneutral, and purple = hydrophilic.

As previously observed with other TRIM family members^31–33^, the asymmetric unit features a single, elongated dimer (Figure 3B). The dimer interface is extensive (5300Å^2^) and is formed primarily by the antiparallel association of the coiled-coil domains, with additional contributions from the RING and second B-box domains. The central coiled-coil domain is comprised of a long arm and a short arm that are separated by a hairpin turn. The long arm is a continuous, extended helix (H1) spanning approximately 170Å, packed against which is the short arm, comprised of two short helices (H2 and H3) separated by a region of extended coil. The coiled-coil region forms a left-handed supercoil at both ends, but the centre is roughly parallel with an interdigitated 4-helical bundle. Extending from the N-terminal end of each coiled-coil domain, the RING and second B-box domains associate closely with the hairpin turn region of their symmetry mate (Figure 3B). The RING and second B-box domains share a similar architecture of a central 3-stranded antiparallel β-sheet, associated with a single short helix and several extended loops. A pair of zinc ions stabilizes each domain, coordinated by cysteine and histidine residues (Figure S4B). Electron densities corresponding to the first B-box domain could not be seen (Figure 3A), nor could the zinc ions bound by this domain be located in anomalous difference maps, suggesting that this domain does not associate with the central core; this interpretation was also supported by limited proteolysis experiments (data not shown).

Similarly, electron density was only observed for a single CUE domain (Figure 3A), which we interpreted to be SMARCAD1 CUE1, based on our data indicating its importance to the SMARCAD1-KAP1 interaction. The SMARCAD1 CUE1 domain is formed of 3 helices and demonstrates considerable structural similarity to the CUE domains of the yeast CUE2p and human gp78 proteins^4,7^ (RMSD of around 1.8Å) (Figure 3C). One distinguishing feature of the SMARCAD1 CUE1 domain, however, is an extended α1 helix that extends for an additional 3 turns (Figure 3C), though the function of this extension is unclear. Reassuringly, our model of the SMARCAD1 CUE1 is largely comparable to the structure of the isolated domain determined independently by NMR^35^.

Our structural model shows a single SMARCAD1 CUE1 domain bound to each end of the KAP1 coiled-coil dimer. The majority of contacts (i.e. approximately 700Å^2^ of a total interface area of around 850Å^2^) are between CUE1 and an exposed surface at the C-terminal end of the long arm of the coiled-coil domain of one KAP1 subunit, but the interface also involves the RING domain of the other KAP1 subunit of the homodimer (Figure 3B and D). The interaction surface on the KAP1 coiled-coil domain is comprised principally of exposed hydrophobic residues, supplemented by a few polar residues at the peripheries. Correspondingly, hydrophobic residues appear to be the primary constituents of the SMARCAD1 CUE1 interaction surface; among these residues are the conserved FP motif (i.e. F169 and P170) and part of the conserved di-leucine motif (i.e. L196) (Figure 3D).

Taken together, the structural model suggests that though the SMARCAD1 CUE1 and KAP1 RBCC domains appear *prima facie* to largely resemble other CUE and TRIM RBCC domains respectively, they nevertheless succeed in associating directly and specifically with each other, mainly on the basis of exposed hydrophobic residues. Remarkably, it also provides clear evidence that the SMARCAD1 CUE1 binds a KAP1 domain that bears no structural resemblance to ubiquitin.

### The α1-α3 Surface of SMARCAD1 CUE1 Mediates a Unique Interaction with the KAP1 RBCC

Next, we compared the interaction interface of SMARCAD1 CUE1-KAP1 RBCC with that of a canonical CUE-ubiquitin interaction. Previous structural studies demonstrated that CUE-ubiquitin interactions rely on a hydrophobic pocket centred on the highly conserved ubiquitin I44 residue being filled by the side chains of the conserved hydrophobic MFP motif in the CUE domain (e.g. M19, F20 and P21 of CUE2p-1); this interaction is then further stabilized by electrostatic interactions around the hydrophobic pocket^4^ (Figures 4B and 4C).

**Figure 4.**
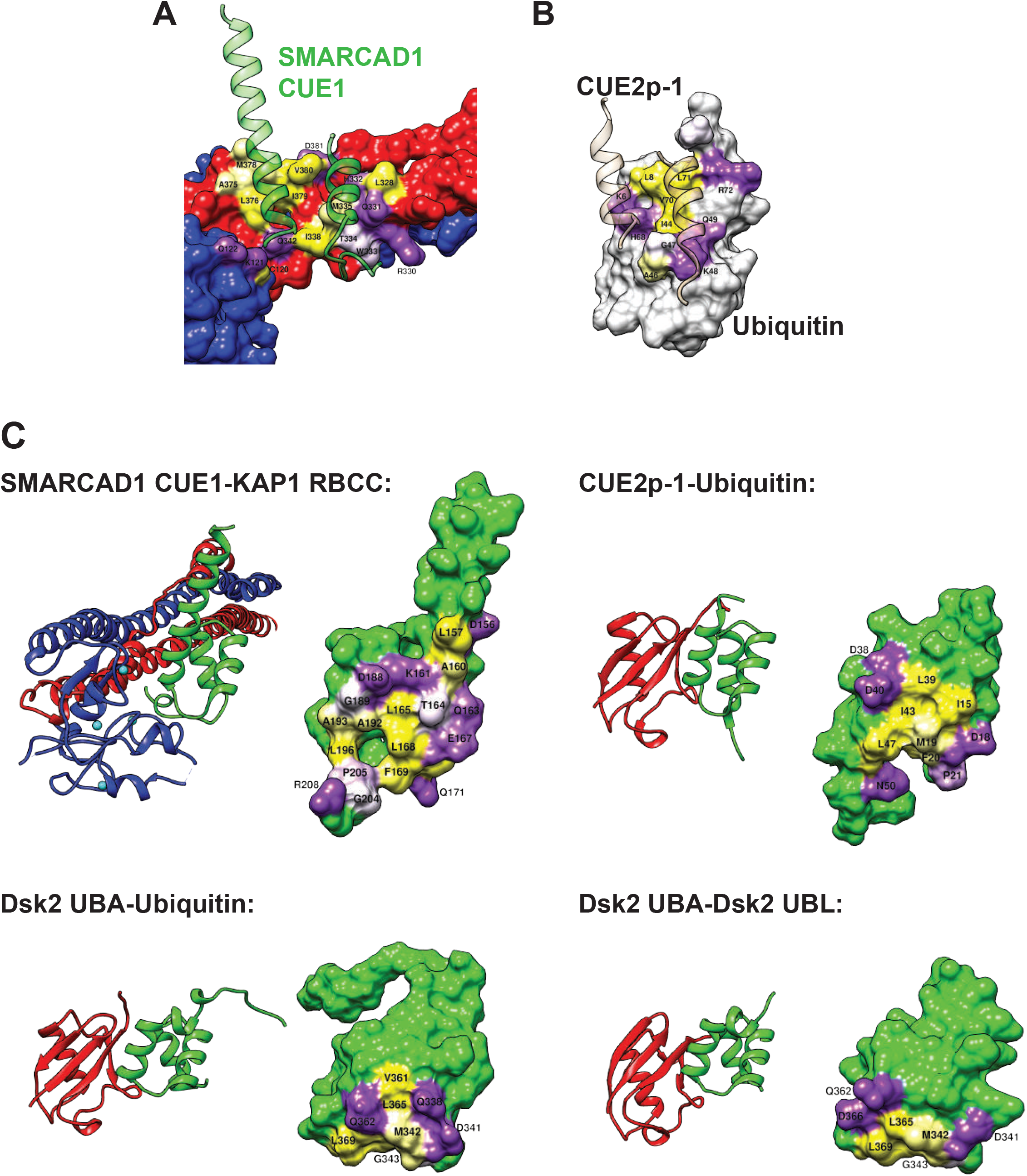
SMARCAD1 CUE1 recognizes KAP1 RBCC with a comparable surface as used for canonical CUE/UBA-Ub binding. **A**. Interaction surface of KAP1, with involved residues coloured according to hydrophobicity as in Figure 3D. The SMARCAD1 CUE1 domain is shown in ribbon format (green), with the α2 helix omitted for clarity. CUE1 recognizes 2 clusters of exposed hydrophobic residues on the KAP1 coiled-coil domain, the first including KAP1 M378 and the second I338. **B**. The structure of ubiquitin in complex with CUE2p, orientated and coloured as in A. CUE domain is depicted as a ribbon (tan), with the α2 helix omitted. A hydrophobic pocket (formed by L8, V70 & I44) surrounded by electrostatic interactions is employed by ubiquitin to interact with the CUE2p-1 domain, a different mode of interaction compared to the KAP1 RBCC-SMARCAD1 CUE1 interaction (compare Figures 4A and B). **C**. The surface employed by SMARCAD1 CUE1 to bind the KAP1 RBCC is similar to that employed by other CUE and UBA domains for canonical ubiquitin or ubiquitin-like (UBL) domain recognition, and features the exposed faces of the α1 and α3 helices. An exposed hydrophobic surface, supplemented by electrostatic interactions at the peripheries, is a common feature of these interactions. The CUE and UBA domains are coloured green, whilst their ligands are red; the interaction surface is coloured by hydrophobicity as previously.

Interestingly, there is no comparable central hydrophobic pocket on the KAP1 RBCC surface that is filled by SMARCAD1 CUE1. Instead, there are 2 clusters of exposed hydrophobic residues, the first including KAP1 M378 and the second I338 (Figure 4A). The extended α1 helix of SMARCAD1 CUE1 lies over these two hydrophobic clusters – the first pocket established by M378 is occupied by SMARCAD1 T164, whereas the second, featuring KAP1 I338, interacts with SMARCAD1 L168 and F169 residues. As L168 and F169 (together with P170) represent the conserved MFP motif in SMARCAD1 CUE1, it appears that I338 of the KAP1 RBCC plays a role akin to that of the crucial I44 residue of ubiquitin. Other interactions between the KAP1 RBCC and SMARCAD1 CUE1 appear to be between the second hydrophobic cluster in KAP1 and the α3 helix of SMARCAD1 CUE1, and electrostatic interactions at the peripheries of the interaction interface, albeit with the caveat that molecular details of these interfaces may not be fully defined by our structural model given its relatively low resolution. Some of these electrostatic interactions involve the RING domain of the other KAP1 subunit, suggesting that the SMARCAD1-KAP1 interaction requires KAP1 to be in its homo-dimeric state (Figures 3D and 4A).

It is notable, however, that the surface employed by SMARCAD1 CUE1 to interact with KAP1 RBCC is comparable to that employed by other CUE (e.g. CUE2p-1) and UBA domains (e.g. Dsk2 UBA) for canonical binding to mono-ubiquitin and ubiquitin-like (UbL) domains (Figure 4C). This interface is formed from the exposed surfaces of the α1 and α3 helices of the CUE or UBA domains. Although variability is observed in the precise residues employed by a specific UBD for ubiquitin recognition, these surfaces are united by their shared possession of a central core of exposed hydrophobic residues encircled by hydrophilic or polar residues (Figure 4C). Thus, this topographical organization is in accord with the hydrophobic pocket (i.e. ubiquitin) or hydrophobic clusters (i.e. KAP1 RBCC) on the interaction surfaces of their cognate ligands.

As the surface on SMARCAD1 CUE1 that would typically be involved in canonical ubiquitin binding is already deployed and optimized for interacting with the KAP1 RBCC, it is unlikely that SMARCAD1 CUE1 can bind simultaneously to KAP1 RBCC and ubiquitin. While non-canonical UBD-ubiquitin/UbL interactions have been reported^36,37^, this is unlikely to be applicable to SMARCAD1 CUE1 as the dissociation constant for it binding mono-ubiquitin has been measured by NMR titration experiments to be roughly 1.8 mM^35^, too weak to represent a biologically meaningful interaction. Together, these data supports a model for the SMARCAD1 CUE1 domain uniquely mediating a distinct interaction with KAP1 rather than functioning as a generic UBD.

Unfortunately, we were unable to determine a motif unique to SMARCAD1 CUE1 to help predict other CUE (or other UBDs) that might similarly recognize a ligand structurally distinct from ubiquitin (data not shown). This is in large part because conserved residues, such as the integral MFP motif, are still present in SMARCAD1 CUE1 and in fact retain important roles in the SMARCAD1 CUE1-KAP1 RBCC interaction; additional residues, for instance T164 (Figure 6A), have simply been co-opted to customize the exposed α1-α3 helix surface for KAP1 RBCC binding. It is thus apparent that sequence conservation is insufficient evidence to implicate a given UBD in canonical ubiquitin binding, and the possibility of it mediating non-canonical interactions with non-ubiquitin(-like) ligands has to be tested empirically.

### The KAP1 Interaction Surface is Not Conserved Amongst TRIM Proteins

The structures of TRIM5α, TRIM20 and TRIM25 have previously been reported^31–33^, and were compared to that of KAP1 (Figure 5A). Although all the TRIM proteins have a similar, central anti-parallel coiled-coil, it is notable that each coiled-coil has a unique geometry, and that the degree to which each helix deviates off its axis varies between the TRIM proteins. Importantly, in contrast to KAP1, none of the other TRIM proteins has the pattern of two clusters of exposed hydrophobic residues on their coiled-coil domains in the region of the coiled-coil where SMARCAD1 CUE1 interacts with KAP1, suggesting that this is not a conserved architectural feature of TRIM proteins (Figure 5A). We conclude that, compared to other proteins of the TRIM family, KAP1 possesses a unique pattern of exposed hydrophobic residues, which form an interaction surface to bind the CUE1 domain of SMARCAD1.

**Figure 5.**
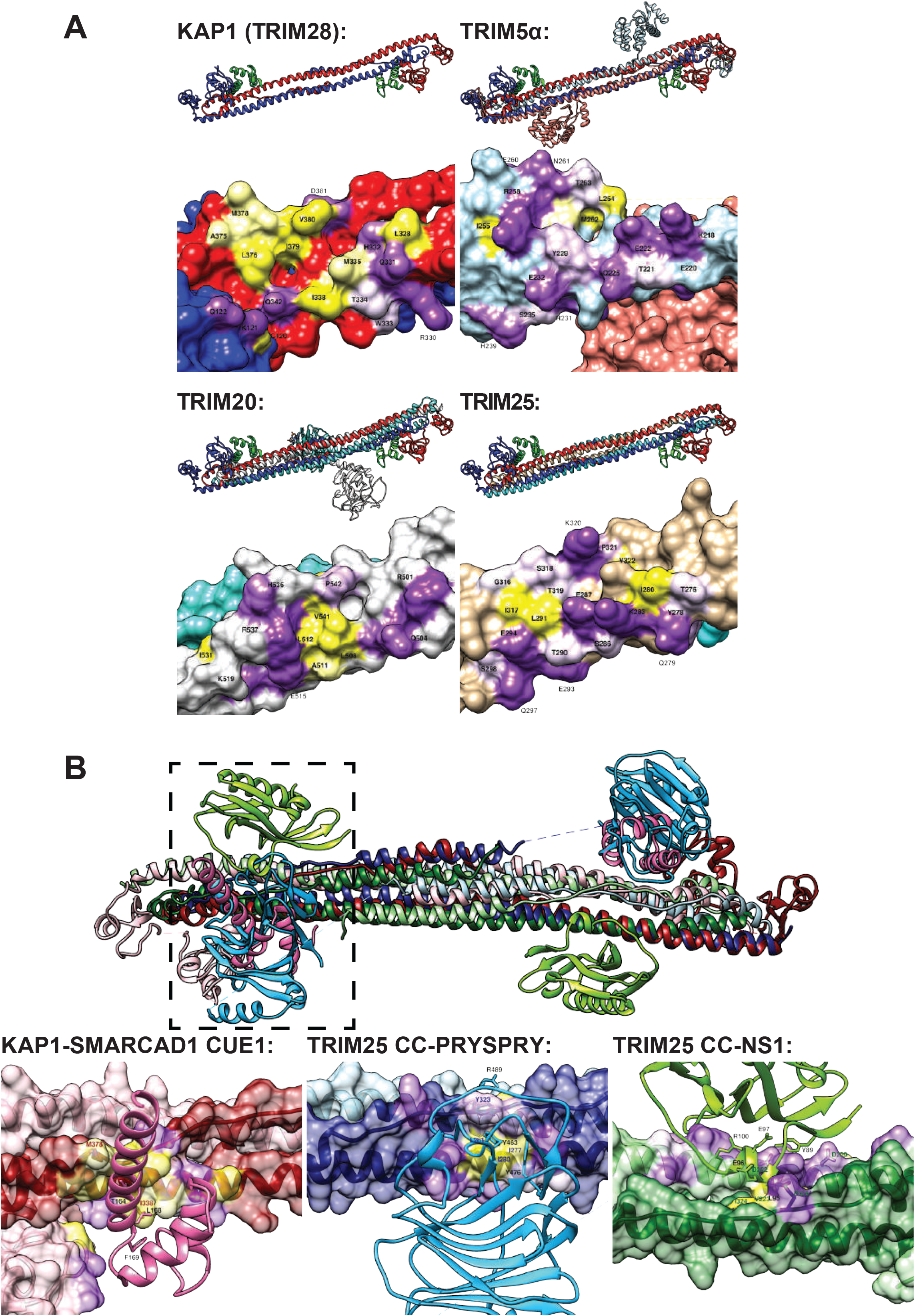
Analysis of the KAP1 interaction surface recognized by SMARCAD1 CUE1. **A**. The structure of the KAP1 (TRIM28) RBCC is similar to that of other TRIM proteins, but the precise geometry of the antiparallel coiled-coil of each TRIM protein is unique, as seen from the misalignment when overlain. Hydrophobicity analysis of equivalent surfaces on different TRIM proteins suggests that the presence of two clusters of exposed hydrophobic residues, used by KAP1 for SMARCAD1 CUE1 binding, is not a conserved structural element. **B**. The KAP1 RBCC-SMARCAD1 CUE1 structure (chains coloured in shades of red) is compared with those of TRIM25 coiled-coil in complex with the TRIM25 PRYSPRY domain (blue) or NS1 (green). The TRIM25 PRYSPRY and SMARCAD1 CUE1 domains bind to the same side, and to similar (but not identical) regions of the coiled-coil domain. In contrast NS1 binds to the opposite side of the coiled-coil domain. The area enclosed by dotted lines is shown magnified in the images at the bottom, with the coiled-coil domains orientated in comparable alignment, and the interaction surface coloured by hydrophobicity.

Rittinger and colleagues have recently elucidated the crystal structure of the coiled-coil domain of TRIM25 in complex with either the TRIM25 PRYSPRY domain or the influenza A non-structural protein 1 (NS1)^38^. Notably, they observe that both the TRIM25 PRYSPRY domain and NS1 protein bind to exterior surfaces of the coiled-coil, albeit on opposite sides of the coil. Comparison of these structures to our SMARCAD1 CUE1-KAP1 RBCC model reveals that all of these interactions localize to a similar region towards the end of the anti-parallel coiled-coil. In fact, it is striking that the region on the TRIM25 coiled-coil surface recognized by the PRYSPRY domain is nearly equivalent to that utilized on the KAP1 coiled-coil domain for SMARCAD1 CUE1 binding (Figure 5B). Nevertheless, the contacts underpinning these distinct interactions are different – the TRIM25 coiled-coil-PRYSPRY interaction being dependent a pair of conserved tyrosine residues (i.e. Y463 and Y476) that project into the coiled-coil^38^, in contrast to SMARCAD1 CUE1, which forms hydrophobic interactions with two clusters of exposed hydrophobic residues on the KAP1 coiled-coil (Figure 5B). While it has not possible to identify sequence features or structural elements that enable prediction of interactions involving the coiled-coil domains of TRIM proteins from this limited data, it is intriguing to speculate that a general feature of TRIM proteins is that the region towards the end of their anti-parallel coiled-coil acts as an interaction surface for varying partner proteins.

### Validation of the Structural Model by Mutagenesis

We sought to validate our structural model by mutation of key residues that might be expected to abrogate the SMARCAD1-KAP1 interaction. Residues identified by the structural model to potentially be important for the interaction were mutated to alanine in the previously described SMARCAD1 CUE1,2 and KAP1 RBCC fragments (Figure 2). Additionally, the 4 residues that were co-targeted in the CUE1mt,2 mutant were also individually mutated. The purified CUE1,2 mutants were tested in binding assays. Notably, SMARCAD1 T164A, L168A or F169A mutation abrogated the ability of the CUE1,2 fragment to bind KAP1 RBCC to the same extent as the quadruple CUE1 mutation (Figure 6A, lanes 5, 8, 9 and 17). Our structural model indicates that SMARCAD1 T164 forms contacts with M378, on which the first hydrophobic cluster of the KAP1 interaction surface is centred, while SMARCAD1 L168 and F169 contact KAP1 I338, a component of the second hydrophobic cluster (Figure 3D, right; Figure 4A).

**Figure 6.**
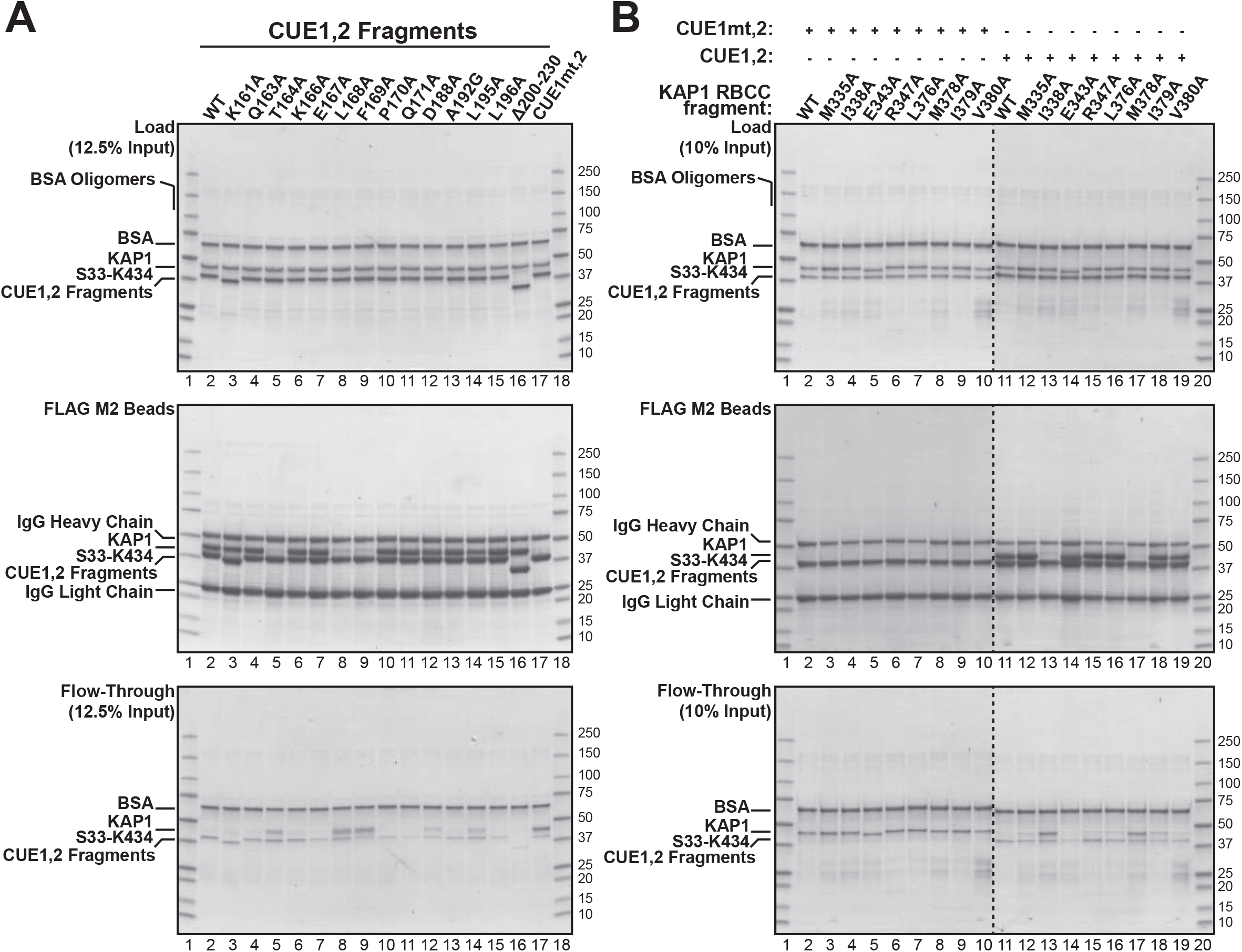
Validation of the KAP1 RBCC-SMARCAD1 CUE1 structure by mutagenesis. **A**. Effect on the SMARCAD1-KAP1 interaction of amino acid changes in the SMARCAD1 CUE1,2 fragment. **B**. As in A, but amino acid substitutions made in KAP1 RBCC (S33-K434).

We next investigated whether mutations targeting these KAP1 residues would also affect SMARCAD1 binding, as predicted by the structural model. Gratifyingly, mutation of KAP1 M378 (first hydrophobic patch; Figure 6B, lane 17) and I338 (second hydrophobic patch; lane 13) completely abrogated binding to SMARCAD1 CUE1,2. Somewhat curiously, despite the prominent effect of the M378A mutation, we noted that other mutations targeting the first hydrophobic patch in KAP1 (i.e. L376A, I379A and V380A) did not perceptibly affect binding (Figure 6B, lanes 16, 18 and 19). Likewise, despite M335 being a constituent part of the second hydrophobic patch, its mutation did not in itself affect binding either (Figure 6B, lane 12). Nevertheless, these results together provide strong empirical support for the notion that the mechanism by which the SMARCAD1 CUE1 domain binds to KAP1 is by recognition of two clusters of exposed hydrophobic residues on the coiled-coil domain.

The CUE1mt,2 fragment used in Figures 1 and 2 is a composite of the F169A, P170A, L195A and L196A mutations, and was designed to target both the conserved MFP and the di-leucine repeat motifs. Strikingly, the inability of the CUE1mt,2 to bind KAP1 appears to be almost exclusively due to the F169A mutation, as neither P170A, L195A nor L196A point mutation individually had a noticeable effect on the ability of the CUE1,2 fragment to bind KAP1 RBCC (Figure 6A, compare lanes 10, 14 and 15 with lane 9 and 17). This helps further support the hypothesis that the SMARCAD1 CUE1-KAP1 RBCC interaction is fundamentally different from canonical CUE-ubiquitin interactions, as the proline of the MFP motif and the second leucine of the di-leucine repeat form important hydrophobic contacts with ubiquitin. Indeed, L47A mutation of the CUE2p-1 domain (analogous to the L196A mutation of SMARCAD1 CUE1) completely disrupts ubiquitin binding *in vitro*^4^.

In addition to the crucial role played by the two exposed hydrophobic clusters in one KAP1 subunit’s coiled-coil domain for binding to a SMARCAD1 CUE1 domain, our model also identified potentially important interactions between the same CUE1 domain and the RING domain of the other KAP1 subunit of the KAP1 homodimer. These interactions involve P170 and Q171 of SMARCAD1 CUE1, and residues C120, K121 and Q122 of the KAP1 RING domain (see Figure 3D, right). However, substitution of P170 or Q171 for alanine had no noticeable effect on the ability of the SMARCAD1 CUE1,2 fragment to bind KAP1 (Figure 6A, lanes 10 and 11). Likewise, mutations targeting other auxiliary interactions outside the hydrophobic clusters (e.g. E167A, D188A, 196A, Δ200-230) were equally ineffective in disrupting the interaction (Figure 6A, lanes 7, 12, 15 and 16). Finally, the KAP1 E343A mutation, designed to target an interaction at the periphery of the interface and outside of the hydrophobic clusters, also had no discernible effect on the interaction (Figure 6B, lane 14). Thus, it appears that these electrostatic interactions are indeed peripheral, in that they may, at least individually, only offer minor contributions to the interaction.

Collectively, the mutagenesis experiments empirically validate the structural model and support the conclusion that the mechanism underpinning the SMARCAD1-KAP1 interaction relies principally upon the SMARCAD1 CUE1 domain recognising two exposed hydrophobic clusters on the KAP1 coiled-coil, with minor contributions from contacts between residues at the periphery of the interaction surface. The first hydrophobic cluster is centred upon KAP1 M378, while the second is crucially reliant upon I338. Notably, this mechanism of interaction is fundamentally different from the filled hydrophobic pocket employed in canonical CUE-ubiquitin interactions. Although the presence of exposed hydrophobic clusters at the end of the coiled-coil domain does not appear to be a shared architectural feature of other TRIM proteins, this may yet transpire to be a conserved mechanism by which other CUE domains form direct protein-protein interactions with their partner proteins.

## Discussion

### CUE Domains Mediate Protein Interactions

CUE domains are generally regarded as protein interfaces for only one ligand, namely ubiquitin (or, much less commonly, ubiquitin-homology domains). Here, however, we show that the first CUE domain of SMARCAD1 – a classical CUE domain – recognizes a ligand structurally distinct from that of ubiquitin, and that it uses this unique property to mediate a stable, direct protein-protein interaction with KAP1. Specifically, SMARCAD1 CUE1 recognizes two exposed clusters of hydrophobic residues situated on the exterior surface of the KAP1 coiled-coil domain. This is a novel mechanism that hitherto has not been associated with CUE domains, or even with UBDs generally. Importantly, it raises the point that when interrogating the function of an uncharacterized UBD, the possibility of it mediating transient or constitutive protein-protein interactions, rather than just interactions with ubiquitylated partner proteins, should also be considered. Its relevance *in vivo* is underscored by the finding that mutations in the SMARCAD1 CUE domain indeed disrupt what is normally a very stable SMARCAD1-KAP1 complex in human cells.

Ubiquitin-UBD interactions are typically weak, with dissociation constants in the hundred-micromolar range, consistent with modest interaction surfaces^4–7^. Indeed, these biochemical properties lend themselves particularly well to transient interactions dependent on the ubiquitylation status of a protein; in turn, this feature of ubiquitin-UBD interactions helps make ubiquitylation such an important and versatile post-translational regulatory mechanism^39^. Nevertheless, it now appears that a proportion of UBD-mediated interactions is independent of ubiquitylation and may even represent more stable interactions. For instance, the yeast Def1 protein uses its CUE domain to specifically recruit the Elongin-Cullin E3 ubiquitin ligase complex to RNA polymerase II via a UbH domain in Ela1^9^, while we have described here how a very stable SMARCAD1-KAP1 complex is achieved in a SMARCAD1 CUE1-dependent fashion.

One inevitable question raised by these observations is the mechanism by which a CUE interaction can be rendered significantly more stable than classical ubiquitin-UBD binding. Although our structural model has not conclusively addressed this issue, it is possible that the avidity of the SMARCAD1 CUE1-KAP1 RBCC interaction is enhanced by the requirement for the SMARCAD1 CUE1 to recognize two, rather than a single, exposed hydrophobic cluster. This is supported by the observation that mutagenesis compromising the integrity of either hydrophobic cluster alone was sufficient to abrogate the interaction. Another consideration is that the modest dissociation constants for ubiquitin-based interactions are typically measured for the interaction between a UBD and mono- (or di-)ubiquitin in isolation. Hence, these reported dissociation constants may give the impression of weaker interactions than in reality, since a weak UBD-ubiqutin interaction, in the presence of additional specificity domains, could result in an overall stable protein-protein interaction between a UBD-containing protein and its ubiquitylated partner^3,39^. Yet, the SMARCAD1-KAP1 interaction is distinctive in being very stable, but apparently dependent only on the CUE1-RBCC interaction, a phenomenon also observed in human cells. Free ubiquitin cannot out-compete recombinant KAP1 RBCC for binding to SMARCAD1 CUE1, and once formed, the SMARCAD1-KAP1 complex withstands dilution or washing with high salt buffers. Overall, our findings help support a model of CUE domains, and presumably UBDs as a whole, potentially fulfilling a general role in mediating protein-protein interactions, besides functioning as ubiquitin-binding domains. Further work will obviously be required to confirm the generalizability of this hypothesis.

### Regulation of SMARCAD1

SMARCAD1 is a poorly characterized, putative chromatin remodeler that is conserved from budding yeast to humans, whose main interaction partner in human cells is KAP1^16,25^. Intriguingly, the *S. cerevisiae* homologue, Fun30, only has a single CUE domain^12^. Tellingly, multiple sequence alignment clusters the Fun30 CUE domain with the *second* SMARCAD1 CUE domain, suggesting that CUE1, which we have shown to be essential for interaction with KAP1, may have been acquired later in evolution. Correspondingly, a KAP1 homologue has not been identified in budding yeast, perhaps suggesting co-evolution of the first SMARCAD1 CUE domain and the SMARCAD1-KAP1 interaction. While our work was being finalized for publication, Mermoud and colleagues reported that mouse SMARCAD1 also interacts with KAP1 via a CUE1-RBCC interaction^40^, nicely supporting the conclusions reported here. No structural data were reported, though their data showed that this interaction is indeed important for recruiting SMARCAD1 to KAP1 target genes in stem cells^40^. Interestingly, complementary results from Morris and colleagues linked DNA repair involving BRCA1-BARD1 to SMARCAD1 through an interaction between human SMARCAD1 and ubiquitylated histone H2A via its CUE domains, though whether CUE2 is solely responsible for this interaction was not reported^22^. Thus, while much remains to be learned about the role of SMARCAD1’s CUE domains in cellular regulation, it is clear that the stable SMARCAD1-KAP1 complex requires the CUE1-RBCC interaction also in human cells.

### The KAP1 RBCC Domain as an Interaction Interface

Our structural model of the SMARCAD1 CUE1-KAP1 RBCC complex confirms that the KAP1 RBCC adopts a structural architecture comparable to other TRIM proteins. It also complements the previously reported structure of the C-terminal KAP1 PHD-bromodomains, meaning that the 3D structure of nearly the entire KAP1 protein has been elucidated, though it will be crucial to determine the organization of the various domains in relation to one another^29^. TRIM proteins adopt an elongated appearance, dominated by a central coiled-coil, and accessorized not only by N-terminal RING and B-box domains at the apices, but also by additional C-terminal protein domains protruding from the centre of the central helix^31–33^. This modular assembly of multiple proteins domains render TRIM proteins particularly adept as scaffold proteins, recruiting the desired molecular machinery to specific cellular or genomic locations, which with reference to KAP1, would presumably be in recruiting epigenetic and DNA repair machinery variously to sites of DNA damage or heterochromatin.

Notably, we show that the SMARCAD1-KAP1 interaction does not occur via a discrete protein domain of KAP1. Rather, the SMARCAD1 CUE1 recognizes an exposed surface, with a particular pattern of hydrophobic residues and orientated in a specific geometry, which is created by homo-dimerization of the KAP1 coiled-coil domains. Intriguingly, Rittinger and colleagues recently reported that a comparable exposed surface of the TRIM25 coiled-coil domain mediates interaction with the TRIM25 PRYSPRY domain, while influenza A NS1 can bind to the opposite side of the TRIM25 coiled-coil domain, though simultaneous interaction of both factors cannot be accommodated due to distortion of the linker connecting the H2 and H3 helices by NS1 binding^38^. It is unclear why the ends of the coiled-coil domains appear to be hotspots for interactions, though it is possible that the close proximity of the flexible linker between the H2 and H3 helices in that region of the coiled-coil allows unique interaction surfaces to be created without disrupting the intermolecular packing of the H1 helices. Nevertheless, protein-protein interactions involving the exterior surface of coiled-coil domains may represent a general feature of TRIM proteins, and it would be intriguing to investigate which other TRIM proteins adopt a similar mode of interaction with their partner proteins. However, we note that it is difficult to predict *a priori* whether a certain partner protein will interact with a specific TRIM protein in this manner, given the considerable exposed areas of each coiled-coil, and the idiosyncrasies in the precise geometry of the helix of each TRIM protein. In spite of these caveats, our findings tentatively support a model of TRIM proteins functioning as an interaction interface by two mechanisms – first, through discrete protein domains that autonomously mediate protein-protein interactions, and second, by supporting interactions that involve exposed surfaces created by oligomerization of the coiled-coil domain.

## Methods

### Plasmids

Human SMARCAD1 and KAP1 cDNA was cloned into a pET28a-SUMO vector, a kind gift from Peter Cherepanov (Crick Institute), for bacterial expression. cDNA was also cloned into the pcDNA4/TO vector (Thermo Fisher Scientific) for use with T-Rex inducible mammalian expression system. Epitope tags, point mutations and truncations were introduced by standard PCR methods. Full length KAP1 and various KAP1 mutants were cloned into the GST expression vector, pGEX6P1 (GE Healthcare). SMARCAD1 CUE1,2 and KAP1 RBCC mutants used to validate our structure were generated by GenScript (Piscataway, USA).

### Generation of Stable Cell Lines

293 T-Rex cells were depleted of endogenous SMARCAD1 by GIPZ lentiviral shRNA (Dharmacon) knockdown, before being rescued with doxycycline-inducible expression of exogenous, shRNA-resistant, FLAG-tagged SMARCAD1 or SMARCAD1 CUE1mt,2mt, using the T-Rex system (ThermoFisher Scientific). Individual colonies were isolated. Doxycycline titration identified concentrations that resulted in exogenous SMARCAD1 being expressed at approximately endogenous levels.

### Preparation of Cell Extracts & Protein Detection

Soluble bacterial extracts were prepared in GST-L-Zn buffer (20mM Tris, 100mM NaCl, 10% (v/v) glycerol, 0.1% (v/v) NP-40, 50μM ZnSO_4_, 5mM β-ME; pH7.90 at 4°C), treated with lysozyme (2mg/mL, Sigma), sonicated in a Bioruptor (Diagenode), and digested with micrococcal nuclease (2000 gel units/mL, NEB). Mammalian whole cell extracts were prepared in Triton lysis buffer (50mM Tris, 150mM NaCl, 1mM EDTA, 1% (v/v) Triton X-100; pH7.50 at RT) supplemented with 1X protease inhibitor cocktail (284ng/mL leupeptin, 1.37μg/mL pepstatin A, 170μg/mL PMSF, 330μg/mL benzamidine, and sonicated in a Bioruptor^®^ (Diagenode). Protein concentrations were determined using the Bradford assay (Bio-Rad Protein Assay) calibrated with a BSA standard curve.

Criterion™ pre-cast XT Bis-Tris 4-12% or TGX™ 5-15% gradient gels (Bio-Rad) were used for SDS-PAGE. Purified proteins were detected by InstantBlu (Expedion), silver (SilverQuest Silver Staining Kit, Invitrogen), or SYPRO^®^ Ruby (ThermoFisher Scientific) staining. Alternatively, Western blotting was performed according to standard techniques using Amersham™ Protran Premium 0.45μm nitrocellulose (GE Healthcare). Membranes were pre-stained with Ponceau S solution (Sigma). The primary antibodies used here were: anti-SMARCAD1 (Bethyl A301-593A) 1:1000, anti-KAP1 (Abcam ab10483) 1:1000, anti-α-tubulin (clone TAT-1) 1:10000, anti-ubiquitin (Enzo Life Sciences clone P4D1) 1:1000, anti-HA (clone 12CA5) 1:10000, and anti-FLAG (Sigma F7425) 1:1000. Either sheep anti-mouse IgG or donkey anti-rabbit IgG HRP-linked F(ab’)_2_ fragments (GE Healthcare) diluted 1:10000 was used as the secondary antibody.

### Immunoprecipitation

FLAG-tagged proteins were immunoprecipitated using 15μL of anti-FLAG^®^ M2 affinity gel (Sigma) (or mouse IgG beads (Sigma) as controls) from cell extract containing 2.5mg of total protein per reaction. After incubation at 4°C for 3 hours, beads were washed thrice in lysis buffer, and eluted by boiling in SDS loading buffer.

### Expression & Purification of Recombinant Proteins

SMARCAD1 and its derivatives (e.g. CUE1,2 and CUE1 fragments) were expressed in BL-21 CodonPlus (DE3)-RIL (Stratagene) *E. coli* cells, KAP1 and its derivatives in BL-21 CodonPlus (DE3)-RP (Stratagene), and the SMARCAD1 CUE1,2-KAP1 RBCC complex was co-expressed in Rosetta2 (DE3) cells (Novagen). Expression was induced with 0.5mM IPTG at either 16°C (full-length SMARCAD1) or 30°C (all other constructs), for either 3 (CUE1,2 fragments) or 6 hours (all other constructs).

SMARCAD1 was nickel-affinity purified with a 5mL HisTrap HP column (GE Healthcare), then dialysed overnight at 4°C against P-100 buffer (10mM sodium phosphate, 100mM NaCl, 10%(v/v) glycerol, 5mM β-ME; pH7.50 at 4°C) in the presence of 100μg of recombinant Ulp1 (a SUMO protease). Subsequent chromatographic steps were a 5mL HiTrap Heparin HP column (GE Healthcare), ProSwift WCX-1S (ThermoFisher Scientific) for SMARCAD1 CUE1,2mt and SMARCAD1 CUE1mt,2mt, and ProSwift SAX-1S (ThermoFisher Scientific). The final fractions were concentrated using a Microcon spin concentrator (Millipore) with a 50K MWCO and exchanged into P-100 buffer.

SMARCAD1 CUE1,2 and CUE1 fragments were first affinity purified using 3mL of Ni-NTA agarose (Qiagen). The SUMO tag was cleaved by recombinant Ulp1 (140μg) during dialysis against Q-100 buffer (10mM Tris, 100mM NaCl, 10%(v/v) glycerol, 5mM β-ME; pH7.90 at 4°C), and depleted by reloading the sample over the 3mL of Ni-NTA resin and collecting the unbound flow-through. If required, these purifications were followed by ion exchange chromatography on a 1mL Mono Q 5/50 GL column (GE Healthcare). The samples were concentrated with Amicon Ultra-4 10K MWCO spin concentrators (Millipore).

As KAP1 and the KAP1 RBCC contain zinc-finger domains, they were expressed whilst cultured in LB supplemented with 50μM ZnSO_4_ or ZnCl_2_, and all buffers used in the purification protocol contained 50μM ZnSO_4_ or ZnCl_2_. KAP1 was purified using a 5mL HisTrap HP column, dialysed against P-100 buffer in the presence of recombinant Ulp1 (100μg), then loaded onto a 5mL HiTrap Heparin HP column. The eluate was concentrated to a volume of approximately 4mL using an Amicon Ultra-15 30K MWCO spin concentrator, before being loaded onto a 120mL HiPrep 16/60 Sephacryl S-400 HR gel filtration column (GE Healthcare). The sample was re-concentrated using a spin concentrator; the final buffer was GF-150Zn buffer (10mM Tris, 150mM NaCl, 50μM ZnSO_4_, 2mM DTT; pH7.90 at 4°C).

KAP1 RBCC (S33-K434) was purified using 3mL of Ni-NTA agarose, cleaved with recombinant Ulp1 (140μg) during dialysis against Q-100 buffer, and depleted of its SUMO tag as described above. This was followed by chromatography on a 1mL Mono Q 5/50 GL column, and peak fractions dialyzed against Q-100 buffer.

The KAP1 RBCC-SMARCAD1 CUE1,2 complex was purified on a 5mL HisTrap HP column, dialyzed against Q-100 buffer, during which the SUMO tag was cleaved by recombinant Ulp1 (140μg), then loaded onto an 8mL Mono Q HR 10/10 column (GE Healthcare). Peak fractions were concentrated to approximately 4mL using an Amicon Ultra-15 30K MWCO spin concentrator, before further purification by gel filtration chromatography using a 120mL HiLoad 16/60 Superdex 200 column (GE Healthcare), the peak of which was re-concentrated using a spin concentrator.

### Reconstitution of SMARCAD1-KAP1 Complex & Analytical Gel Filtration

180μg of purified FLAG-SMARCAD1 was mixed with 180μg of purified HA-KAP1 in SK reconstitution buffer (10mM sodium phosphate, 200mM NaCl, 10% (v/v) glycerol, 0.1% (v/v) NP-40; pH7.50 at RT) and 5mM β-ME. The complex was then reconstituted by sequential affinity purification with anti-HA (3F10) affinity matrix (Roche) and anti-FLAG M2 affinity gel (Sigma). Bound proteins were eluted respectively with HA (1mg/mL) and FLAG peptide (500μg/mL, Peptide Chemistry core facility, Francis Crick Institute), prepared in P-100 buffer.

Analytical gel filtration chromatography was performed using a 4×300mm MAbPac SEC-1 column (ThermoFisher Scientific). 375ng of protein was loaded per run, and eluted isocratically in P-200 GF buffer (10mM sodium phosphate, 200mM NaCl, 2mM DTT; pH7.50 at 4°C).

### Crystallization, Structure Determination & Refinement

For crystallization an N-terminally SUMO-tagged KAP1 G53-K434 fragment was overexpressed in *E.coli* BL21(DE3)-R3-pRARE2 cells. Cells were grown at 37°C in TB medium supplemented with 50μg/mL kanamycin until an optical density of 2-3, then induced with 0.3 mM IPTG and incubated overnight at 18°C. Purification was as described above for KAP1 S33-K434, with the exception of the final purification step of size exclusion chromatography using a HiLoad 16/60 Superdex S200 column, where a buffer containing 50mM Hepes pH7.5, 500mM NaCl, 5% glycerol and 0.5mM Tris (2-carboxyethyl) phosphine (TCEP) was used instead. KAP1 G53-K434 was concentrated to 10mg/ml using a Millipore 30,000 MWCO centrifugal concentrator and mixed with SMARCAD CUE1,2 (purified as described above) in a 1:1.1 ratio (slight excess of SMARCAD CUE1,2). Crystallization was performed by sitting drop vapour diffusion and crystals were grown from conditions containing 1.2M sodium malonate, 0.5% Jeffamine ED-2003 and 0.1M HEPES pH7.0, with a 1:2 protein to precipitant drop ratio. Crystals were loop mounted and transferred to a cryoprotectant solution comprising the well solution supplemented with 25% ethylene glycol, before being flash-cooled in liquid nitrogen.

A SAD dataset extending to 5.5Å was collected at Diamond Light Source beamline i03 and the data were processed using DIALS^41^. The structure was solved using Phenix autosol^42^ using the intrinsic anomalous signal of the zinc ions, and the initial phases were improved substantially by solvent flattening. Model building was performed using either existing crystal structures of fragments or template derived models, which were directly fitted in to the experimentally phased maps based on zinc ions (i.e. RING and B-box domains) or recognizable secondary structure elements (i.e. coiled-coil and CUE1 domain). Side chain positions were chosen from preferred rotamers that minimized clashes with neighbouring atoms and the structure was refined using Phenix refine^43^, using both NCS and reference model restraints, and a single B-factor per residue. A summary of the data collection and refinement statistics is shown in Table 1.

**Table 1.**
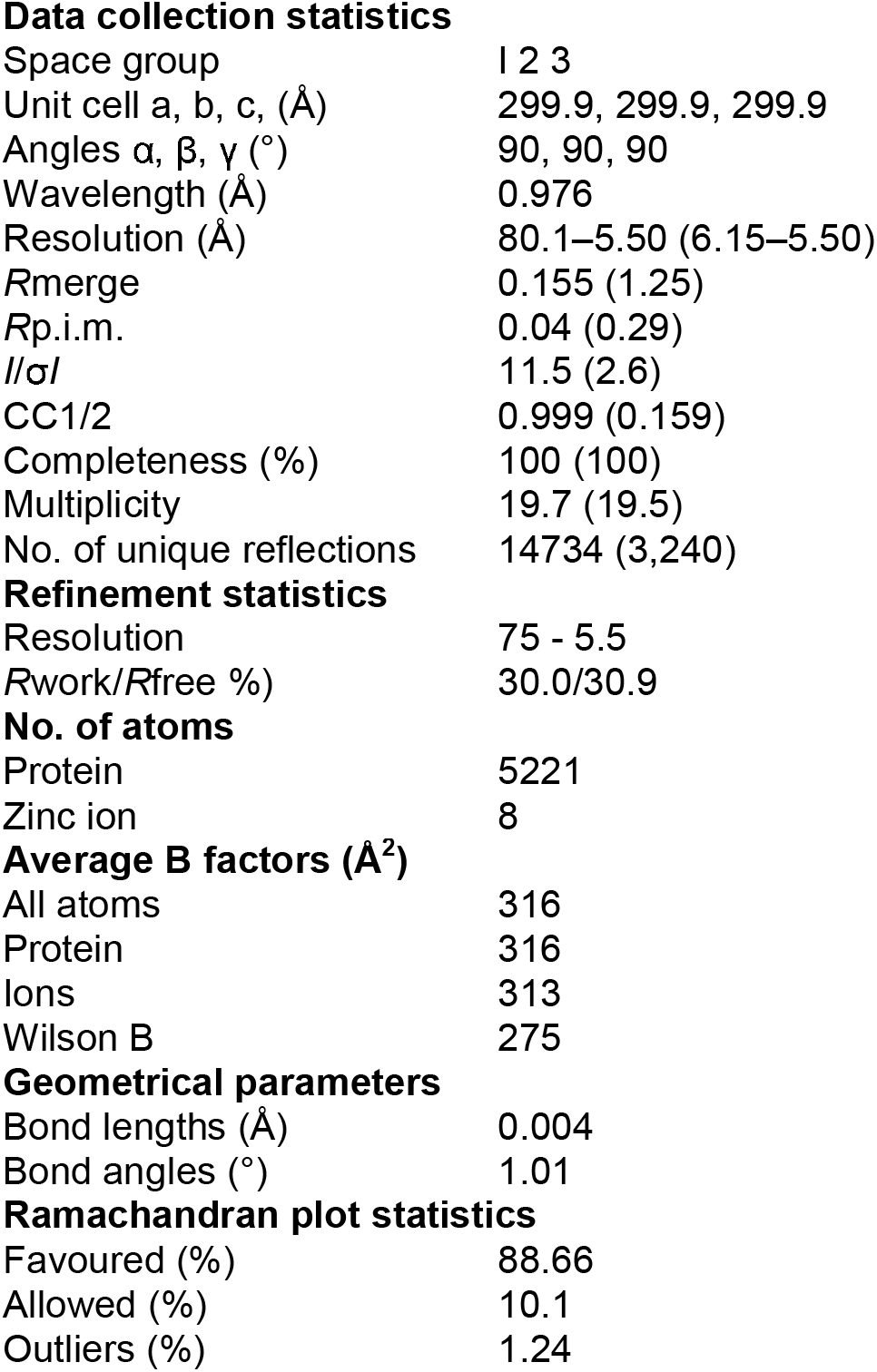
Data collection and refinement statistics for the KAP1 RBCC-SMARCAD1 CUE1,2 complex

|  |  |
| --- | --- |
| Space group | I 2 3 |
| Unit cell a, b, c, (Å) | 299.9, 299.9, 299.9 |
| Angles $\alpha$, $\beta$, $\gamma$ (°) | 90, 90, 90 |
| Wavelength (Å) | 0.976 |
| Resolution (Å) | 80.1–5.50 (6.15–5.50) |
| *R*<sub>merge</sub> | 0.155 (1.25) |
| *R*<sub>p.i.m.</sub> | 0.04 (0.29) |
| *I*/σ*I* | 11.5 (2.6) |
| CC1/2 | 0.999 (0.159) |
| Completeness (%) | 100 (100) |
| Multiplicity | 19.7 (19.5) |
| No. of unique reflections | 14734 (3,240) |
| **Refinement statistics** |  |
| Resolution | 75 - 5.5 |
| *R*<sub>work</sub>/*R*<sub>free</sub> %) | 30.0/30.9 |
| **No. of atoms** |  |
| Protein | 5221 |
| Zinc ion | 8 |
| **Average B factors (Å<sup>2</sup>)** |  |
| All atoms | 316 |
| Protein | 316 |
| Ions | 313 |
| Wilson B | 275 |
| **Geometrical parameters** |  |
| Bond lengths (Å) | 0.004 |
| Bond angles (°) | 1.01 |
| **Ramachandran plot statistics** |  |
| Favoured (%) | 88.66 |
| Allowed (%) | 10.1 |
| Outliers (%) | 1.24 |

Refined structures were visualized and analysed using UCSF Chimera^44^. For comparative analysis, atomic coordinates were obtained from the PDB using accessions 1OTR (ubiquitin-CUE2-1 complex)^4^, 1WR1 (Dsk2 UBA-ubiquitin complex)^45^, 2BWE (Dsk2 UBA-Dsk2 UBL complex)^46^, 4TN3 (TRIM5α)^32^, 4CG4 (TRIM20)^33^, 4LTB (TRIM25)^31^, 6FLN (TRIM25 coiled-coil-TRIM25 PRYSPRY complex)^38^ and 5NT2 (TRIM25 coiled-coil-NS1 complex)^38^.

### SMARCAD1-KAP1 Binding Assays

For the binding assay with purified full-length SMARCAD1 and KAP1, 7μg of each was mixed together in a 280μL binding reaction containing SK binding buffer (10mM Tris pH7.50 at RT, 150mM NaCl, 10% (v/v) glycerol, 0.01% (v/v) NP-40, 50μM ZnSO_4_), 0.1mg/mL BSA, and 5mM β-ME. The reactions were adjusted to a final sodium chloride concentration of 200mM. The binding reactions were incubated at 4°C for 1 hour before being immunoprecipitated overnight at 4°C with 20μL of anti-FLAG M2 affinity gel (Sigma) per reaction. The beads were washed in 500μL of SK-200 buffer (10mM Tris pH7.50 at RT, 200mM NaCl, 10% (v/v) glycerol, 0.01% (v/v) NP-40, 50μM ZnSO_4_, 5mM β-ME) three times before the beads were eluted with 30μL of 2X SDS-PAGE loading buffer by heating the samples to 100°C for 5 minutes.

The binding assays involving purified fragments of SMARCAD1 (i.e. CUE1,2 and CUE1) and KAP1 (i.e. RBCC) were performed similarly, with the following slight adjustments: 9.6μg of each protein was used in a 240μL binding reaction incubated for 1 hour at 4°C, and immunoprecipitated with 15μL of anti-FLAG M2 affinity gel for 3 hours at 4°C before elution as described above. The effect of ubiquitin on the SMARCAD1-KAP1 interaction was investigated by adding purified recombinant, monomeric ubiquitin (Boston Biochem) to the binding reaction.

The affinity resin of immobilized SMARCAD1 CUE1,2 fragment was prepared by saturating the binding capacity of the M2 resin with three-fold as much purified FLAG-tagged protein (approximately 12.2nmol protein/mL resin), incubating the beads at 4°C overnight, before washing off unbound protein. To 20μL of CUE1,2-coupled resin, *E. coli* extracts containing GST-tagged KAP1 fragments (2.5mg of total protein per reaction) were added and incubated at 4°C for 3 hours. The beads were washed thrice in SK-200 buffer thrice, before being eluted as above.

### Limited Tryptic Proteolysis

Limited tryptic digestion was performed in trypsin buffer (20mM Tris pH7.40, 50mM NaCl, 1mM CaCl_2_, 2mM DTT) using 1/1000 (w/w) the amount of trypsin as purified protein. The reactions were stopped by addition of a protease inhibitor cocktail. For Edman degradation, digested samples were resolved by SDS-PAGE, transferred onto an Amersham Hybond P 0.45 PVDF membrane (GE Healthcare), and stained with Ponceau S (Sigma), following which, selected bands were excised. Edman degradation (5 cycles each) was performed by AltaBioscience.

For intact molecular weight mass spectrometry, the digested samples were first incubated with 50mM DTT to remove β-ME adducts (from the purification buffers). Tryptic peptides were removed using an Ultrafree-CL centrifugal filter unit with a 5K MWCO (Millipore). LC/MS grade formic acid (Fisher Scientific) was added for a concentration of at least 0.2% (v/v) and pH.

## Data Accessibility

Atomic coordinates and structure factors for the X-ray structure of the KAP1 RBCC-SMARCAD1 CUE1,2 complex have been deposited in the PDB with accession 6H3A. Other data and constructs used in this study are available from the corresponding author upon reasonable request.

## Acknowledgements

This work was supported by the Francis Crick Institute (which receives its core funding from Cancer Research UK (FC001166), the UK Medical Research Council (FC001166), and the Wellcome Trust (FC001166)), and by a grant from the European Research Council, Agreements 693327 (TRANSDAM). ML was supported by a UCL Overseas Research Scholarship. The Proteomics, Peptide Chemistry, and Cell Services Science Technology Platforms of the FCI provided us with expert assistance and service. We thank Andreas Ehrensberger, Marcus Wilson, Valerie Pye and Peter Cherepanov (FCI) for their support and time spent on this project, and Katrin Rittinger, Kay Hofmann and Ivan Dikic for their insightful comments on the manuscript.

## Author Contributions

ML and JQS conceived of the project and designed the experiments. ML made bacterial expression constructs, purified the recombinant proteins, and performed all binding experiments. JN solved the structure, and HA purified proteins for crystallography. HW made the human cell lines expressing SMARCAD1. JQS and OG supervised the work, ML and JQS wrote the paper, and all the authors helped edit it.

## Supplementary Figure Legends

**Figure S1.**
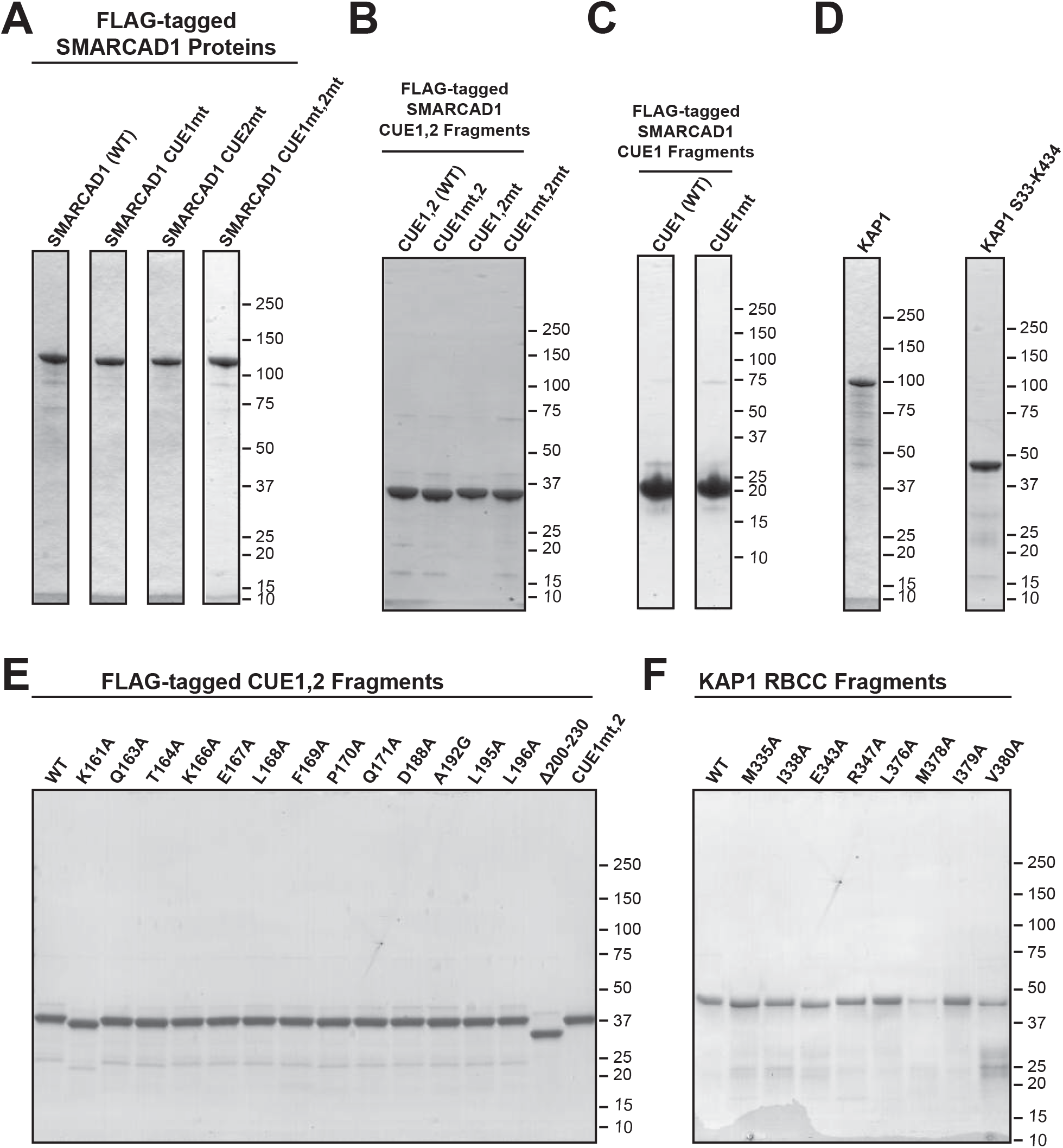
Purified proteins used in this study. **A**. Full-length SMARCAD1, wild type or mutant versions, expressed in *E. coli* and purified. **B**. Wild type and mutant CUE1,2 fragments (S95-N347), expressed in *E. coli* and purified. **C**. Wild type and mutant CUE1 fragments (S95-E237), expressed in *E. coli* and purified. **D**. Purified, recombinant full length KAP1 and the minimal soluble KAP1 S33-K434 fragment. **E**. FLAG-tagged SMARCAD1 CUE1,2 fragments used to validate the structural model. **F**. As in E, but KAP1 RBCC fragments.

**Figure S2.**
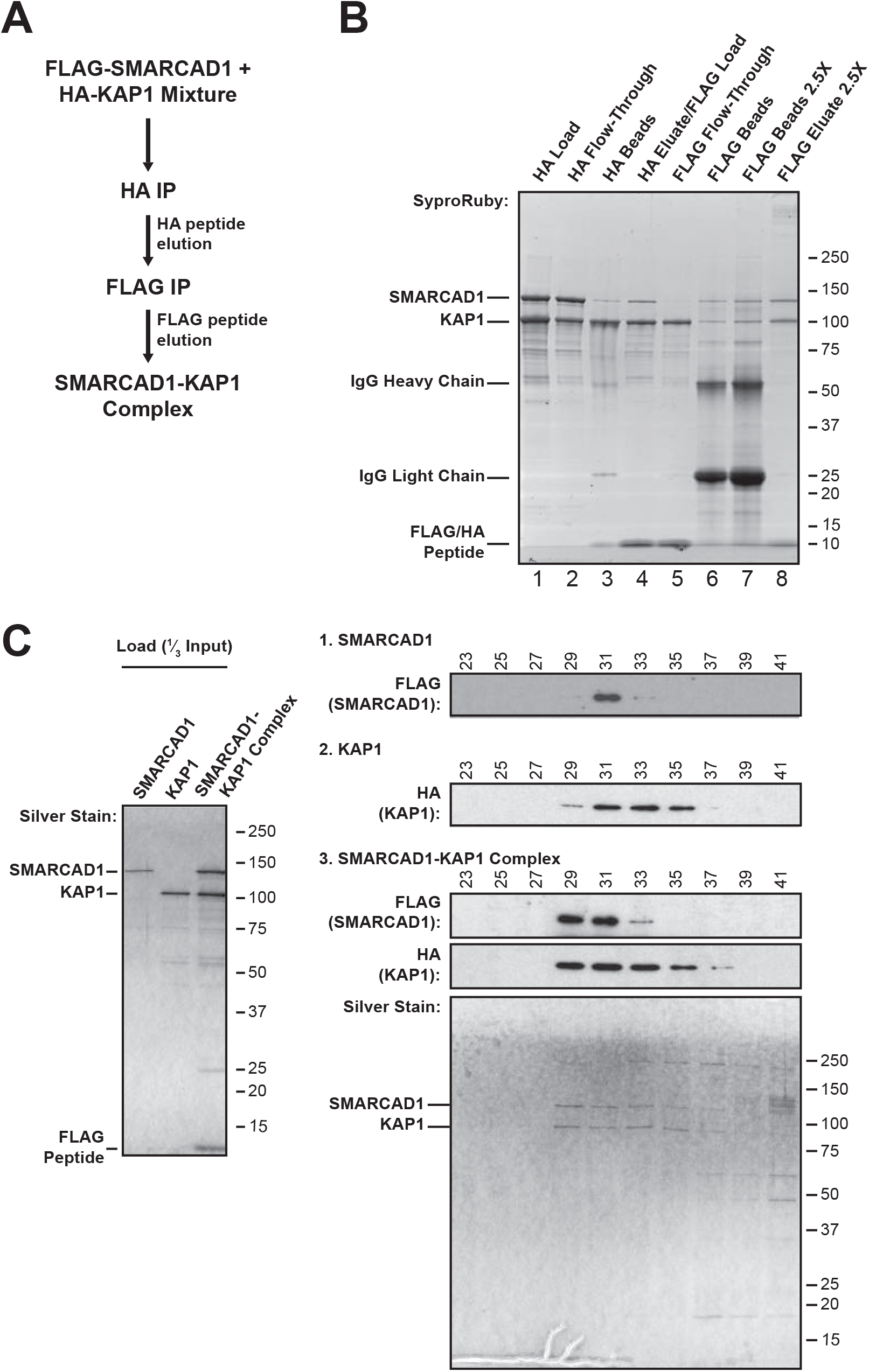
Reconstitution of the SMARCAD1-KAP1 complex. **A**. Diagram of sequential affinity purifications performed to reconstitute the SMARCAD1-KAP1 complex *in vitro*. **B**. Representative gel of the SMARCAD1-KAP1 complex reconstitution protocol. The final reconstituted SMARCAD1-KAP1 complex is pure and comprised of near-stoichiometric quantities of each protein. **C**. Analytical gel filtration chromatography confirming that SMARCAD1 (1.) and KAP1 (2.) elute as single peaks, while SMARCAD1 and KAP1 of the reconstituted SMARCAD1-KAP1 complex (3.) precisely co-elute in earlier eluting fractions.

**Figure S3.**
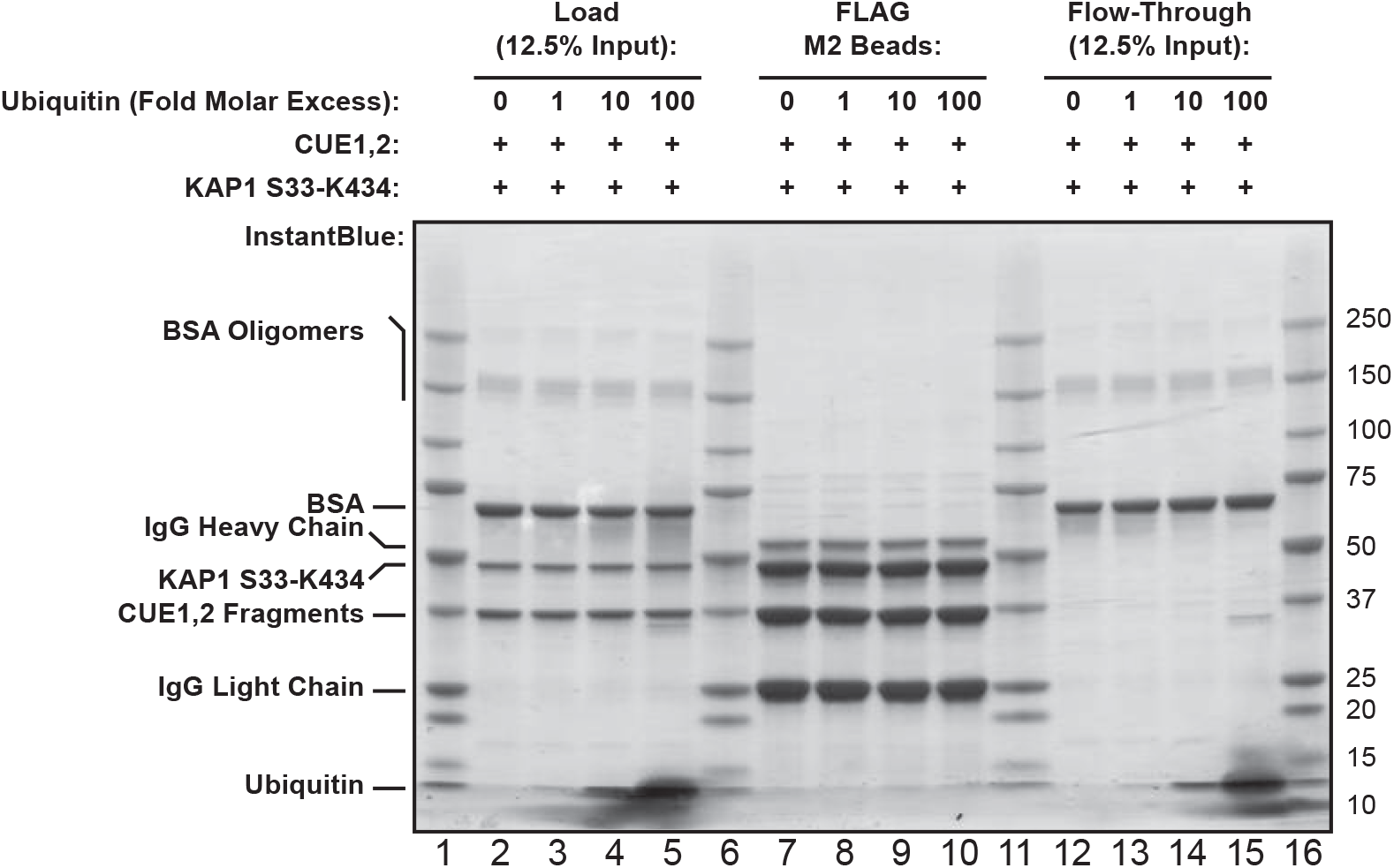
KAP1 RBCC has greater affinity for the SMARCAD1 CUE domains than mono-ubiquitin. A large molar excess of ubiquitin fails to outcompete binding of the KAP1 S33-K434 fragment to immobilized purified CUE1,2.

**Figure S4.**
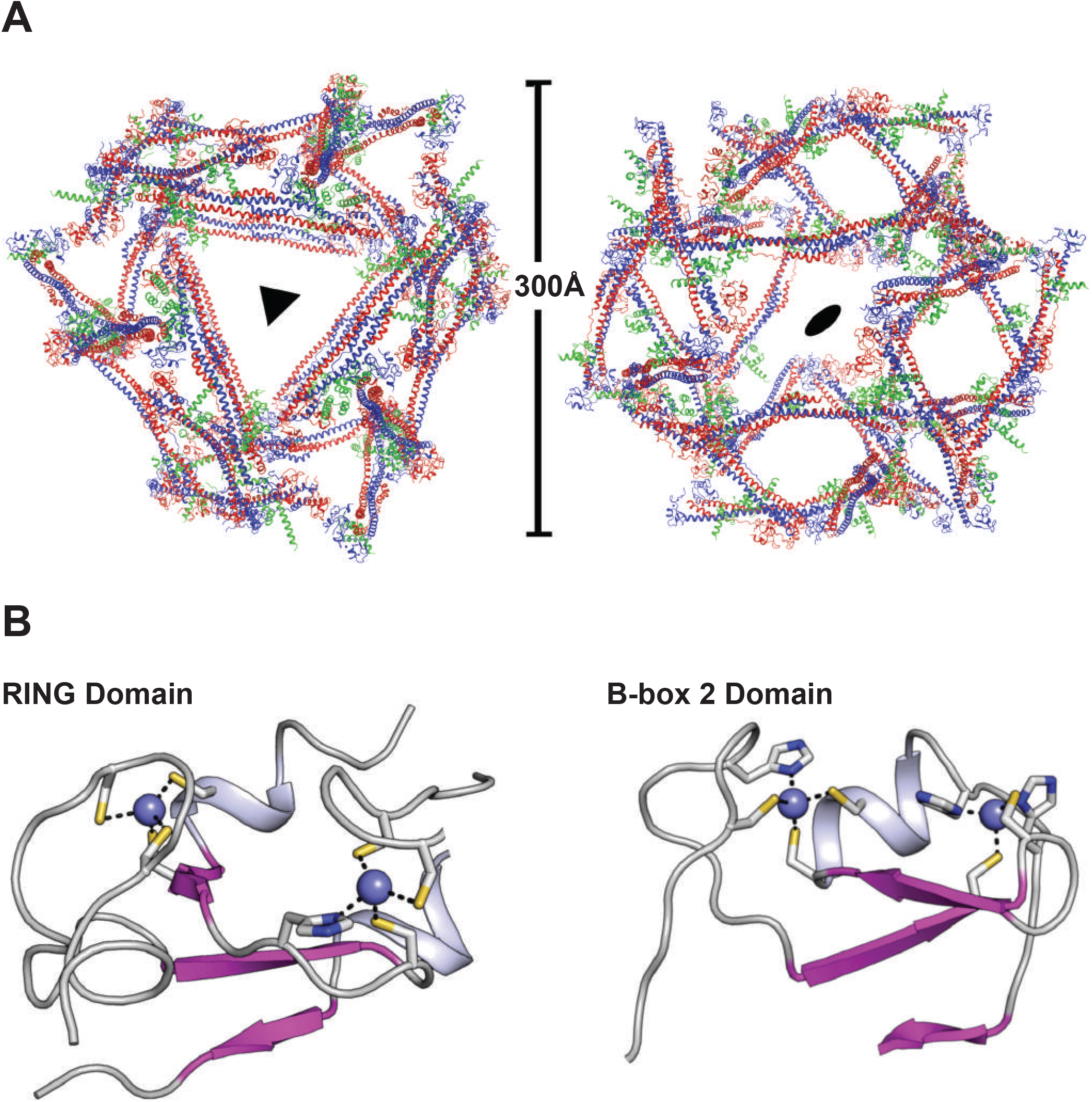
Additional features of the KAP1 RBCC-SMARCAD1 CUE1,2 crystal structure. **A**. The unit cell is a proteinaceous cage with internal voids, explaining the extremely high solvent content of the crystal. **B**. The RING and second B-box domains of KAP1 are compact domains consisting of a central 3-stranded antiparallel β-sheet, short helix and several extended loops, and each is coordinated by 2 zinc ions.

